# Distribution and survival strategies of diazotrophs in the Arctic Ocean revealed by global-scale metagenomic analysis

**DOI:** 10.1101/2022.10.28.514325

**Authors:** Takuhei Shiozaki, Yosuke Nishimura, Susumu Yoshizawa, Hideto Takami, Koji Hamasaki, Amane Fujiwara, Shigeto Nishino, Naomi Harada

## Abstract

Nitrogen fixation is the major source of reactive nitrogen in the ocean and has been considered to occur specifically in low-latitude oligotrophic oceans. Recent studies have shown that nitrogen fixation also occurs in the polar regions and thus is a global process, although the physiological and ecological characteristics of polar diazotrophs are not yet known. Here, we successfully reconstructed genomes, including that of cyanobacterium UCYN-A (*Candidatus* ‘Atelocyanobacterium thalassa’), from metagenome data corresponding to 111 samples isolated from the Arctic Ocean. These diazotrophs were highly abundant in the Arctic Ocean (max., 1.28% of the total microbial community), suggesting that they have important roles in the Arctic ecosystem and biogeochemical cycles. Diazotrophs in the Arctic Ocean were either Arctic-specific or universal species. Arctic-specific diazotrophs, including Arctic UCYN-A, had unique gene sets (e.g., aromatics degradation) and/or a very small cell size (<0.2 µm), suggesting adaptations to Arctic-specific conditions. Universal diazotrophs were generally heterotrophs and commonly had the gene that encodes the cold-inducible RNA chaperone, which presumably makes their survival possible even in deep, cold waters and polar regions. Thus both types of diazotroph have physiological traits adaptable to their environments, which allow nitrogen fixation on a global scale.

## INTRODUCTION

Nitrogen fixation is the process by which specialized prokaryotes (diazotrophs) convert dinitrogen gas to ammonia. It has long been thought that nitrogen fixation occurs mainly in the N-depleted tropical and subtropical regions where cyanobacterial diazotrophs are prevalent ^1^. However, recent studies have demonstrated that nitrogen fixation occurs even in the N-rich polar regions ^2-5^.

In the Arctic Ocean, sea ice exists throughout the year, and the seawater temperature remains below 5°C. The Arctic surface layer is relatively stable despite the high latitude due to the presence of low-salinity water, and some regions become oligotrophic in summer ^6,7^. During the autumn and winter, mixing processes become active due to the frequent strong wind and atmospheric cooling, and nitrogenous nutrients are supplied to the surface layer throughout the Arctic Ocean ^7^. Furthermore, underwater irradiance varies dramatically throughout the year, with solar radiation disappearing in winter. The Arctic environment is largely different from that in tropical and subtropical oligotrophic regions, in which the seawater temperature in the mixed layer is maintained above 25°C throughout the year, the thick N-depleted surface layer is rarely disturbed due to the stable water structure, and underwater irradiance is also stable throughout the year ^8^. Given this environmental difference, diazotrophs could have unique strategies for adapting to the Arctic environment.

Information on diazotrophs in the Arctic Ocean is currently limited to the *nifH* sequence. *nifH* encodes the iron protein subunit of nitrogenase, which has a highly conserved sequence that can be used for species identification ^9^. Many *nifH* sequences retrieved from the Arctic Ocean differ from those at lower latitudes ^2,3^,10, suggesting the presence of Arctic-specific diazotrophs. However, *nifH* alone does not explain how diazotrophs have adapted to the Arctic environment. Furthermore, *nifH* alone is not necessarily an indicator of microbial nitrogen fixation ^11,12^. Mise et al. ^12^ recently demonstrated that ∼20% of bacterial genomes with *nifH* in the public database do not have *nifD* and/or *nifK*, which encode essential subunits of nitrogenase, indicating that these genomes represent non-diazotrophs. Therefore, current understandings of Arctic diazotrophs based only on *nifH* information are incomplete.

The recent development of culture-independent genome reconstruction techniques such as metagenome-assembled genomes (MAGs) has changed our understanding of microbial ecology, including that of diazotrophs. Genome-based approaches have been applied in tropical and subtropical studies, revealing previously unknown diazotrophs and their physiology and ecology ^13,14^. Recently, we have newly built a marine MAG catalog, which contains >50,000 genomes derived from 8,466 prokaryotic species that were derived from various marine oceanic regions including the Arctic Ocean ^15^. Here, we explored the genome catalog for Arctic diazotrophic species using *nifH* sequences as an initial marker gene. Notably, we successfully retrieved the genome of symbiotic cyanobacterial diazotroph UCYN-A (*Candidatus* ‘Atelocyanobacterium thalassa’) from metagenomic data of the Arctic Ocean. UCYN-A is one of the major diazotrophs in subtropical regions ^16^ and is currently divided into six subclades based on its *nifH* sequences ^17^, of which UCYN**-**A1 and **-**A2 occur exclusively in and fix nitrogen in the Arctic Ocean ^3,4^. The *nifH* sequences of UCYN-A1 and -A2 in the Arctic Ocean are identical to those in the subtropical Ocean ^4^. However, genomic information on UCYN-A in the Arctic Ocean has not been revealed, and thus there are no clues for adaptation mechanisms of UCYN-A to the Arctic environment. We examined the distribution patterns using a global metagenomic database and characterized diazotrophs in the Arctic Ocean by comparative genomic analyses.

## RESULTS AND DISCUSSION

### Overall characteristics of MAGs containing *nifH* that were retrieved from the Arctic Ocean

Metagenomic data derived from 111 samples collected in the Arctic Ocean yielded 6816 prokaryotic MAGs, which include 1095 species according to the OceanDNA MAGs catalog ^15^. Of those MAGs from the Arctic Ocean, *nifH* was detected in nine MAGs (Table 1 and Supplementary Data 1). Four of the nine (Arc-UCYN-A2, Arc-Campylo, Arc-Alpha, and Arc-Gamma-01) were detected with the existing *nifH* universal primers, but the remaining five could not be detected with previous PCR approaches (Table 1 and Supplementary Data 2).

**Table 1.** Summary of *nifH* detected MAGs in the Arctic Ocean

| Genome | Length<br>(Mb) | Genome<br>completeness | *nifH* primer<br>compatibility | *nifH*<br>Cluster | *nifDK, nifENB* | Distribution<br>pattern | Taxonomy |
| --- | --- | --- | --- | --- | --- | --- | --- |
| Arc-UCYN-A2 | 1.48 | 74.3 (99.3)* | ○ | I | ○ | Arctic-specific | Cyanobacteria; *Atelocyanobacterium thalassa* |
| Arc-Bactero | 4.55 | 97.3 | × nifH4 | III | ○ | Arctic-specific | Bacteroidota; genus *Sunxiuqinia* |
| Arc-Campylo | 2.92 | 96.8 | ○ | I | ○ | Arctic-specific | Campylobacterota; genus *Arcobacter* |
| Arc-Alpha | 3.18 | 97.7 | ○ | I | ○ | Universal | Proteobacteria; genus *Novosphingobium* |
| Arc-Gamma-01 | 3.21 | 97.6 | ○ | I | No *nifK* | Arctic-specific | Proteobacteria; genus *Immundisolibacter* |
| Arc-Gamma-02 | 4.14 | 93.9 | × nifH4 | I | ○ | Arctic-specific | Proteobacteria; genus *Psychromonas* |
| Arc-Gamma-03 | 3.74 | 96.4 | × nifH4 | I | ○ | Arctic-specific | Proteobacteria; genus *Oceanobacter* |
| Arc-Gamma-04 | 3.77 | 91.8 | × nifH4 | I | No *nifB* | - | Proteobacteria; genus *Motiliproteus* |
| Arc-Myxo | 3.77 | 84.6 | × nifH1,2,3 | III | No *nifDK*, or *nifENB* | - | Myxococcota; family JABWCM01 |

Genome completeness was estimated by CheckM <sup>57</sup>. \*Calculated by regarding the complete genome of the UCYN-A2 (CPSB-1, Supplementary Data 4) as 100.

Diazotrophs must have at least *nifDK* and *nifEBN* in addition to *nifH* for nitrogen fixation ^11^. Of the nine MAGs, Arc-Gamma-01, Arc-Gamma-04, and Arc-Myxo lacked *nifK, nifB*, and *nifDK* and *nifENB*, respectively, suggesting that these may not be genomes of diazotrophs. However, upon careful inspection, Arc-Gamma-01 had a contig that included *nifD* at the end of the contig. Because *nifHDK* generally form an operon structure in the genome, this MAG may seem to lack *nifK* due to the limitations of a fragmented genome assembly. As Arc-Gamma-01 was categorized into the genus *Immundisolibacter* and the same genus genome was reconstructed with a different method in which all *nif* genes were isolated from Arctic metagenomic data (https://anvio.org/blog/targeted-binning/), we assumed that Arc-Gamma-01 is a diazotroph. However, Arc-Gamma-04 and Arc-Myxo, neither of which is likely to be a diazotroph, were excluded from the subsequent analysis.

Among the Arctic diazotroph MAGs, Arc-Bactero belongs to Cluster III and the rest belong to Cluster I based on the *nifH* sequence ^9^ (Fig. 1a and Table 1). These MAGs were taxonomically classified to Cyanobacteria, Alphaproteobacteria, Gammaproteobacteria, Campylobacterota (formerly categorized as Epsilonproteobacteria), and Bacteroidota at the phylum or class level using the Genome Taxonomy Database ^18^. Diazotrophs belonging to the phylum Bacteroidota have not been reported thus far from the marine water column. The cyanobacterial MAG (Arc-UCYN-A2) was classified to UCYN-A belonging to the subclade UCYN-A2. The remaining six Arctic diazotrophs may represent new species within genera *Arcobacter, Sunxiuqinia, Psychromonas, Oceanobacter, Immunodisolibacter, Novosphingobium* (Table1 and Supplementary Data 1). Thus far, *nifH* of *Arcobacter* has been found in oceans around the globe including the Arctic Ocean ^10,19^,20, and *nifH* of *Novosphingobium* was recently detected in deep waters in the subtropical ocean ^21^. The diazotrophs within genera *Sunxiuqinia, Psychromonas, Oceanobacter*, and *Immunodisolibacter* were newly discovered in this study.

**Fig. 1.**
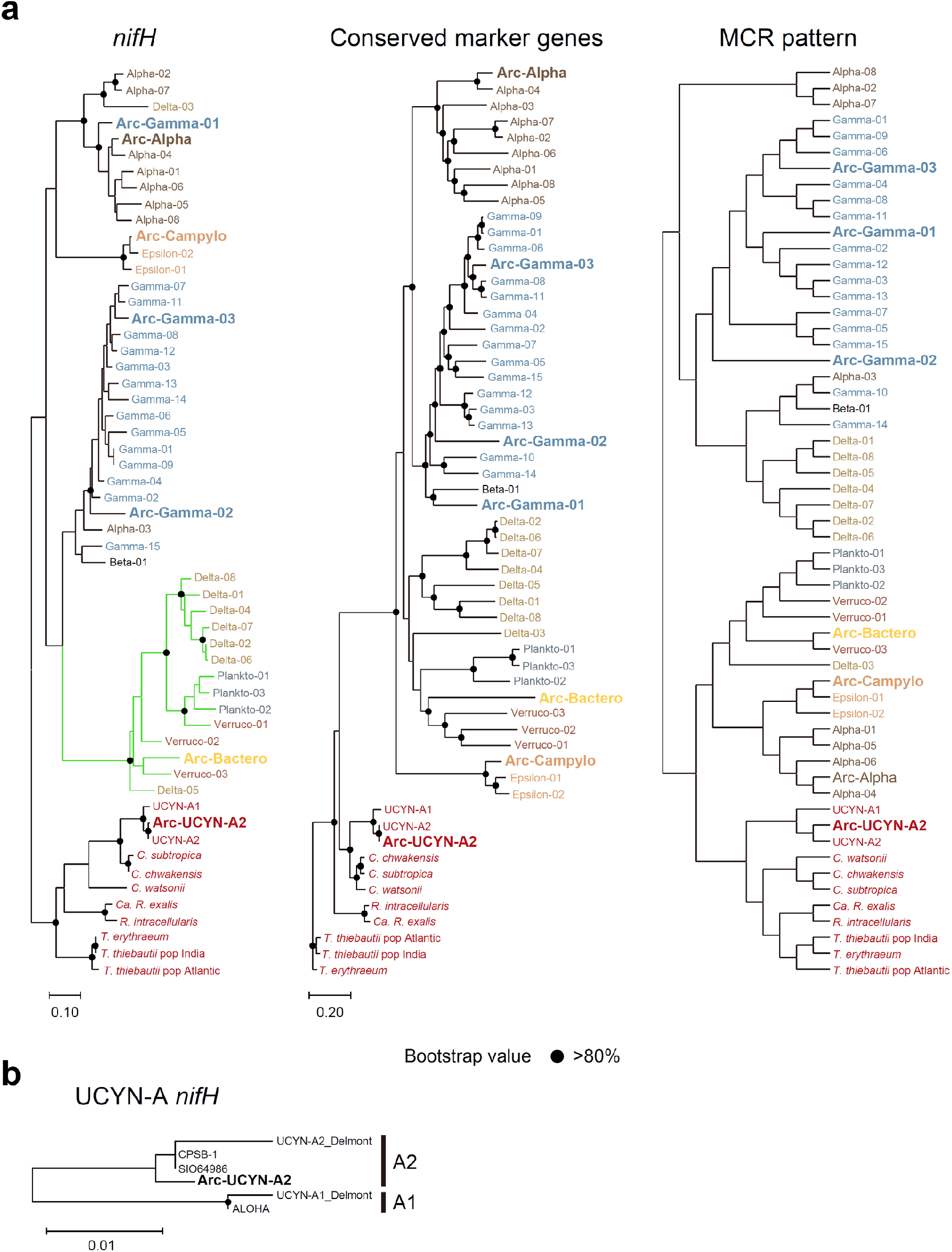
Phylogeny and functional module clustering of diazotrophs. a. three types of dendrograms representing *nifH* phylogeny, conserved marker gene phylogeny, and hierarchical clustering based on the functional module completion ratio (MCR) ^64^. The seven Arctic MAGs (shown in bold) and cyanobacterial diazotrophs *C. chwakensis* and *C. subtropica* are included, along with MAGs constructed from a metagenomic dataset from samples collected in the subtropical ocean ^13,14^. Label colors represent phylum- or class-level taxonomy. The *nifH* phylogenetic tree is rooted with cyanobacterial sequences to be consistent with the other trees. b. *nifH* phylogenetic tree of UCYN-A, for which nearly the whole genome has been assembled (Supplementary Data 4). The phylogenetic tree was estimated with the maximum-likelihood method based on the full length of the *nifH* sequence and the conserved marker genes using PhyloPhlAn. The branches in the *nifH* phylogenetic trees belonging to Clusters I and III are shown with black and green lines, respectively.

The phylogenetic tree based on concatenated sequences of a set of marker genes conserved in all MAGs formed various clusters categorized into every phylum or class in the case of Proteobacteria, as expected, and this result supported the correctness of the phylogenetic assignment for each MAG (Fig. 1a). In contrast, the clusters on the phylogenetic tree based on *nifH* genes did not necessarily reflect the phylogenetic placement because the genes assigned to different taxonomic groups were nested in one cluster. For example, Arc-Gamma-01, which is categorized as belonging to Gammaproteobacteria, was buried within the cluster composed of Alphaproteobacteria, which does suggest that the *nifH* gene in Arc-Gamma-01 was acquired by horizontal gene transfer. In addition to the phylogenetic analysis, we also carried out hierarchical clustering analysis of MAGs based on the pattern of completion ratios of functional modules as calculated by the Genomaple™ system ^22^ (Fig. 1a), which results in the functional classification of the genomes ^23^. Although most clusters consisted of a single taxonomic group, some were made up of phylogenetically distant groups such as Planctomyces, Verrucomicrobia, and Bacteroidota. This indicated that these particular MAGs possess a similar physiological and metabolic potential regardless of their phylogenetic relationships. In addition, as Arctic diazotroph MAGs did not form their own cluster, they did not possess common functional traits that can distinguish them from those from lower latitudes.

### Global distribution of Arctic diazotroph MAGs

We examined the relative abundance of each MAG in a global metagenome database (Fig. S1a) ^15^. Samples were assessed based on size fractionation—total (**≥**0.2 µm; although the viral fraction was not included, this sample is referred to as “total” for convenience), bacterial (0.2–3 µm), and viral (<0.2 µm) fractions—and water column depth—shallow (**≤**200 m), intermediate (200–1000 m), and deep (**≥**1000 m) layers. In general, water temperature at the surface varied widely across the sampled oceanic regions, but in the deep sea it was uniformly low in all regions (Fig. S1b). Similarly, in the metagenome database, water temperatures in the deep layers showed less variation and lower temperatures (median, 2.02 °C) than in other layers (Fig. S1c). The genome abundance of Arctic diazotroph MAGs varied greatly across different size fractions and regions (Fig. 2). In the total fraction (>0.2 µm), the abundance of Arc-UCYN-A2, Arc-Bactero, Arc-Alpha, and Arc-Gamma-01 was high (maximum length-normalized count per million microbe genomes of a total community [CPMM]: 479, 355, 3213, and 12841, respectively) especially in the Arctic Ocean. The maximum CPMM of Arc-Gamma-01 means that the genome abundance reached 1.28% of the total microbial community, which was the highest on a global scale (Fig. 2 and S2a), suggesting that it could be crucial to the Arctic ecosystem and biogeochemical cycles. The abundance of Arc-Alpha also increased in deep water (**≥**1000 m) in low latitudes. In contrast, in the bacterial fraction (0.2–3 µm), most arctic diazotroph MAGs were rarely found. This size-dependent difference in the abundance could be due to the cell size of these bacteria. For example, UCYN-A is a symbiotic diazotroph with haptophytes, and its size including its host is >3 µm ^4,24^. Therefore, UCYN-A was detected in the total fraction but was rarely detected in the bacterial fraction. Similarly, Arc-Bactero, Arc-Alpha, and Arc-Gamma-01 are also likely to have cell sizes >3 µm.

**Fig. 2.**
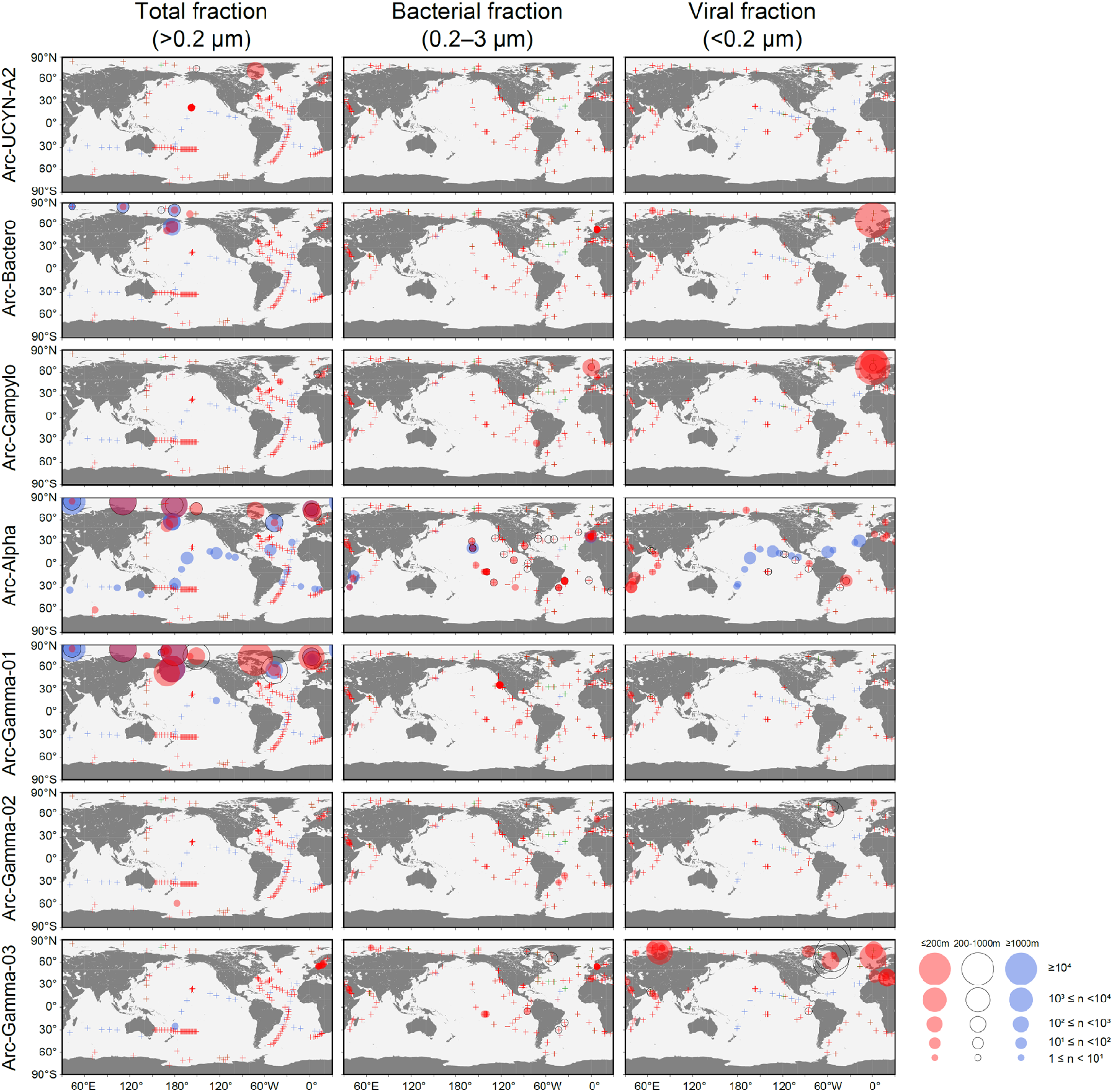
Depth-resolved abundance of Arctic diazotroph MAGs across the global oceans. The abundance data are divided into three size fractions and three depths. The genome abundance was calculated as a length-normalized CPMM in each fraction. The area of each circle is proportional to its indicated abundance. The plus signs indicate the location with CPMM lower than 1.

In the viral fraction (<0.2 µm), the relative abundance of Arc-Bactero, Arc-Campylo, Arc-Gamma-02, and Arc-Gamma-03 was particularly high in the Arctic Ocean samples (maximum CPMM: 14291, 62203, 2530, and 152821, respectively). The CPMM value indicates that Arc-Gamma-03 accounted for up to 15.2% of the viral fraction community. Arc-Bactero was also abundant in the total fraction, which may be attributable to its cell shape (i.e., a slender filamentous cell). Meanwhile, as Arc-Campylo, Arc-Gamma-02, and Arc-Gamma-03 were rarely found in the other fractions, their cell size is likely to be very small, and thus they would have passed through a 0.2-µm filter ^25^. Interestingly, some *nifH* sequences were recently reported to be more abundant in the viral fraction of the Arctic Ocean ^10^. Of the two *nifH* sequences observed in the viral fraction, one was identical to the *nifH* sequence of Arc-Campylo, thus confirming our result.

We further examined the abundance of diazotroph genomes obtained at lower latitudes (Fig. S2). Cyanobacterial diazotrophs were abundant mainly at low latitudes in the total and bacterial fractions as shown in previous microscopy and *nifH*- and genome-based studies ^10,14^,26. The exception was *C. chwakensis*, which was widely found in the Arctic Ocean. Interestingly, some heterotrophic diazotrophs were as abundant or more abundant in the Arctic Ocean than in low latitudes. For example, the relative abundance of Alpha-02 and -04 became high in the Arctic Ocean (maximum CPMM: 324 and 843, respectively) in the total fractions. These heterotrophic diazotrophs also occurred in deep water at low latitudes, and this distribution pattern was similar to that of Arc-Alpha. Among the known diazotrophs, no species was found to have a high relative abundance in only the viral fraction, as was noted for the Arctic diazotrophs Arc-Campylo, Arc-Gamma-02, and Arc-Gamma-03.

In summary, we noted two general distribution patterns of diazotrophs that exist in the Arctic Ocean: (1) diazotrophs that occur almost exclusively in the Arctic Ocean (Arc-Bactero; Arc-Campylo; and Arc-Gamma-01, -02, and -03) and (2) diazotrophs that occur both in the Arctic Ocean and at lower latitudes (Arc-Alpha, *C. chwakensis*, Alpha-02 and -04, and so on) (Table 1 and S1). Based on the size-fractionation data, we were able to estimate the cell size of the Arctic diazotrophs. Arc-UCYN-A2, Arc-Bactero, Arc-Alpha, and Arc-Gamma-01 may have a large cell size of >3 µm, and Arc-Campylo and Arc-Gamma-01 and - 02 are likely to have a very small cell size of <0.2 µm.

### What characteristics allow diazotrophs to occur in the Arctic Ocean?

Arctic-specific microbes other than diazotrophs have previously been described based on 16S rRNA genes and MAGs ^27-29^. Royo-Llonch et al. ^29^ recently investigated the genomic characteristics of Arctic-specific microbes through a large-scale comparison of MAGs from the Arctic Ocean and lower latitudes and showed that Arctic-specific microbes generally have larger genome sizes and shorter minimum generation times, implying a copiotrophic lifestyle. The genome size (1.4–4.5 Mb, Table 1) and minimum generation time (1.0–12 hours, Supplementary Data 1), which was estimated using Growpred ^30^, of Arctic diazotrophs were not significantly different from those of low-latitude microbes in Royo-Llonch et al. ^29^, suggesting no trend of a copiotrophic lifestyle. This may not be surprising for diazotrophs because they can thrive in an oligotrophic environment.

Other possible factors that may make microbes more likely to inhabit the Arctic Ocean are their psychrophilic (grow optimally at <15 °C) or psychrotolerant (survive below the freezing point but grow optimally at 20–25 °C) characteristics ^28,31^. Indeed, *Psychromonas* (Arc-Gamma-02) is a representative of the psychrophilic bacteria, many species of which have been isolated from low-temperature marine environments including polar regions and the deep sea (e.g., ^32,33^). Microbes in cold environments generally have specialized proteins that function alone or in combination to adapt to their growing conditions ^31,34,35^. We examined the genes encoding cold-inducible proteins ^34^ in each MAG (Supplementary Data 1), but we note that some of these cold-inducible proteins are not found only in cold-adapted microbes but can also be present in thermophilic or mesophilic microbes (e.g., ^36^). Among these proteins, the cold-inducible RNA chaperone (*CspA*), which is a protein involved in maintaining RNA structure at low temperatures, is generally shared among microbes in cold environments ^31,34^,35. *cspA* gene was found in all Arctic diazotroph MAGs except that of Arc-UCYN-A2. Furthermore, among the low-latitude diazotrophs examined here, most of the heterotrophic diazotrophs had *cspA* (Supplementary Data 1). Obviously, various genes in addition to *cspA* are involved in cold-environment adaptation, but these results suggested that most marine heterotrophic diazotrophs have the potential to adapt to cold environments.

Heterotrophs can occur from the surface to the deep sea, and thus it is not surprising that heterotrophic diazotrophs inhabiting mainly low latitudes also have *cspA* to adapt to the cold environment of the deep sea (median, 2.02 °C in our dataset). In contrast, cyanobacterial diazotrophs do not have *cspA* except for *C. chwakensis* and *C. subtropica*. Cyanobacteria can grow only in the euphotic layer (<200 m), as they need to perform photosynthesis. In addition, cyanobacterial diazotrophs occurs mainly in high-temperature (>20 °C), low-latitude waters ^1,26^ and thus do not need to have *cspA. C. chwakensis* was found in high abundance in the Arctic Ocean, suggesting that it acquired *cspA* to adapt to the cold environment. Although *C. subtropica* was rarely found in the Arctic Ocean, it is phylogenetically very close to *C. chwakensis* (Fig. 1a and ^37^) and may also have *cspA*. The presence of *cspA* in the genome can explain distribution patterns of diazotrophs that are present both in the Arctic Ocean and at lower latitudes; these include not only *C. chwakensis* but also Arc-Alpha and Alpha-02 and -04. However, *cspA* alone does not explain the occurrence of Arctic-specific diazotrophs. There were no genes encoding cold-inducible proteins that were specific to Arctic diazotroph MAGs with the exception of *ipxP* of Arc-Gamma-02 (Supplementary Data 1), which encodes a protein responsible for lipid A synthesis under low temperatures ^34^. Therefore, in addition to cold-environment adaptations, there are likely to be other reasons for the occurrence of Arctic-specific diazotrophs.

Environmental uniqueness may also be related to characteristics of Arctic diazotrophs. The Arctic Ocean has sea ice throughout the year and is prone to being stratified despite the cold environment due to the fresh water input from melting sea ice and rivers. Arctic rivers also supply a large amount of terrestrial materials to the ocean. Interestingly, Arc-Gamma-01 has a large number of glycosyltransferase genes (57 genes) as compared with low-latitude gammaproteobacterial diazotrophs (14–43 genes) (Supplementary Data 1). Glycosyltransferase is involved in polysaccharide production ^38^. *Crocosphaera* and *Trichodesmium*, which also have a high number of members of this gene family (Supplementary Data 1), produce extracellular polysaccharides (EPSs) and form aggregates ^39,40^ in which EPSs assist with adherence ^38^. Similarly, Arc-Gamma-01 can produce EPSs and may form aggregates and/or attach itself to sea ice. Another unique feature of Arctic diazotroph MAGs is their potential for aromatics degradation. The degradation function of carbazole by Arc-Gamma-01 and of salicylate by Arc-Gamma-03 were not found in any other diazotrophs (Supplementary Data 3). In addition, Arc-Gamma-02 has genes associated with benzene degradation. Colatriano et al. ^41^ recently showed that Chloroflexi, a major bacterial phylum in the Arctic Ocean, may have acquired aromatics degradation genes horizontally from terrestrial bacteria and was subsequently able to grow using material of terrestrial origin. The same thing could have happened to the Arctic diazotrophs. Collectively, the Arctic-specific diazotrophs may have expanded their metabolic potential to adapt to the unique environment of the Arctic Ocean.

It should be note that some Arctic diazotrophs (Arc-Campylo and Arc-Gamma-02 and - 03) are expected to have a very small cell size (<0.2 µm), as was also inferred from an *nifH*-based study ^10^. One characteristic of very small bacteria is a reduced genome size (<2 Mb) ^25^. The genome size of Arc-Gamma-03 (3.7 Mb) is smaller than that of the isolated species in the genus *Oceanobacter* (4.5 and 5.1 Mb), but it is not particularly small. Those of Arc-Gamma-02 (4.1 Mb) and Arc-Campylo (2.9 Mb) are within the range of the genus *Psychromonas* (3.9–5.5 Mb) and *Arcobacter* (2.2–3.2 Mb). Therefore, these Arctic diazotrophs with small cell sizes do not have small genomes. In contrast, the cell size of the Arctic diazotrophs is presumably smaller than that of isolated non-diazotroph species ^32,33^,42-45. In general, bacteria in the field tend to have smaller cell sizes than do cultured strains due to nutrient limitations and predation ^46^. Given that variations in cell size were noticed but appreciable variations in genome size were not, it is likely that environmental stress reduced the cell size of the Arctic diazotrophs. In contrast, the abundance of low-latitude diazotrophs was not notably high in the viral fraction alone (Fig. S2), suggesting that such very small diazotrophs rarely occur at low latitudes, which may mean that Arctic diazotrophs have adapted to greater environmental stress.

### Comparative genome analysis of UCYN-A

We obtained a nearly complete genome of UCYN-A2 from the Arctic Ocean. The *nifH* phylogenetic tree using full-length *nifH* showed that the *nifH* sequence of Arc-UCYN-A2 is distinct from that of low-latitude UCYN-A2 (Fig. 1b), which contradicts the conclusion of a previous study ^4^. This is because the previous study examined only a short PCR-amplified region of the gene. Indeed, the *nifH* sequence targeted by PCR of Arc-UCYN-A2 is identical to that of the low-latitude one.

We further performed a comparative genomic analysis using existing UCYN-A genomes (Supplementary Data 4 and Fig. S3). As noted previously ^47^, substantial differences were found between the UCYN-A1 and -A2 genomes (Fig. S3). In contrast, the intraclade difference was very small; the Arc-UCYN-A2 genome was almost identical to that of UCYN-A2 at low latitudes (ANI 99.42–99.79% as compared with low-latitude UCYN-A2). However, we found that Arc-UCYN-A2 had a *gph* gene, which is used in DNA repair ^48^, that was absent from the low-latitude UCYN-A2. No differences were found except for this *gph* gene when comparing Arc-UCYN-A2 with the low-latitude UCYN-A2 genome. Regarding the *gph* gene, cold-adapted microbes tend to have more genes to repair DNA damage caused by reactive oxygen species, which generally increase inside the cell in cold environments ^34^. Although Arc-UCYN-A2 does not have the *cspA* gene, the *gph* gene in Arc-UCYN-A2 may be used for another strategy for adaptation to the cold Arctic environment.

### *nifH* gene abundance among diazotroph MAGs isolated from Arctic Ocean samples

We quantified the *nifH* copy number of MAGs that showed a particularly high genome abundance in the total fraction of Arctic Ocean and of Arc-UCYN-A2 using qPCR technique. We collected the samples in the Pacific side of the Arctic Ocean in summer for three years. The observed sample sites were located mainly in open-water areas. The sea surface temperature ranged from −1.2 to 7.6 °C. Although nitrogenous nutrients were sporadically high (>1 µM) in the surface water of the Bering Strait, they were generally depleted (< 0.1 µM) in the north of 70°N. Of the targeted diazotrophs, Arc-UCYN-A2, Arc-Alpha, and Alpha-04 were detected in samples collected during each cruise, indicating that these diazotrophs were indeed present in the Arctic Ocean. In contrast, Arc-Bactero, Arc-Gamma-01, *C. chwakensis*, and Alpha-02 were not detected in samples from either cruise. This absence might be related to their habitat area and season, because samples were collected only in the Pacific side open-water area in summer. For example, Arc-Gamma-01 could produce EPSs as mentioned above and thus could attach to the sea ice, which would preclude its sampling from open-water samples. The *nifH* of *C. chwakensis* (former genus *Cyanothece*) was detected in the Atlantic side of the Arctic Ocean ^49^.

UCYN-A2 was found in samples from most of the stations during each cruise and was distributed vertically (Fig. 3a). The maximum *nifH* abundance of UCYN-A2 was 2.9 × 10^3^, 5.8 × 10^5^, and 8.0 × 10^6^ copies L^−1^ in 2015, 2016, and 2017, respectively, and was found near the surface. Arc-Alpha and Alpha-04 were widely distributed in the study region but were detected only at the surface. The maximum for Arc-Alpha (3.3 × 10^2^ copies L^−1^) and Alpha 04 (3.0 × 10^2^ copies L^−1^) was significantly lower than that of UCYN-A2 (*p* < 0.05). We examined the relationship between *nifH* abundance and environmental parameters and found that UCYN-A2 had a significant positive correlation with temperature (*p* < 0.05), although other diazotrophs had no significant relationship with any environmental parameters (*p* > 0.05). This relationship between UCYN-A abundance and temperature was also reported at low latitudes, although the temperature range (22–30 °C) differed from that in the Arctic Ocean (–0.4–7.6 °C) ^50,51^. Hence, the controlling factor seems to be the same even if the clade is different. The positive correlation with temperature indicated that UCYN-A2 can expand its habitat range and should be able to increase in the Arctic Ocean with future warming.

**Fig. 3.**
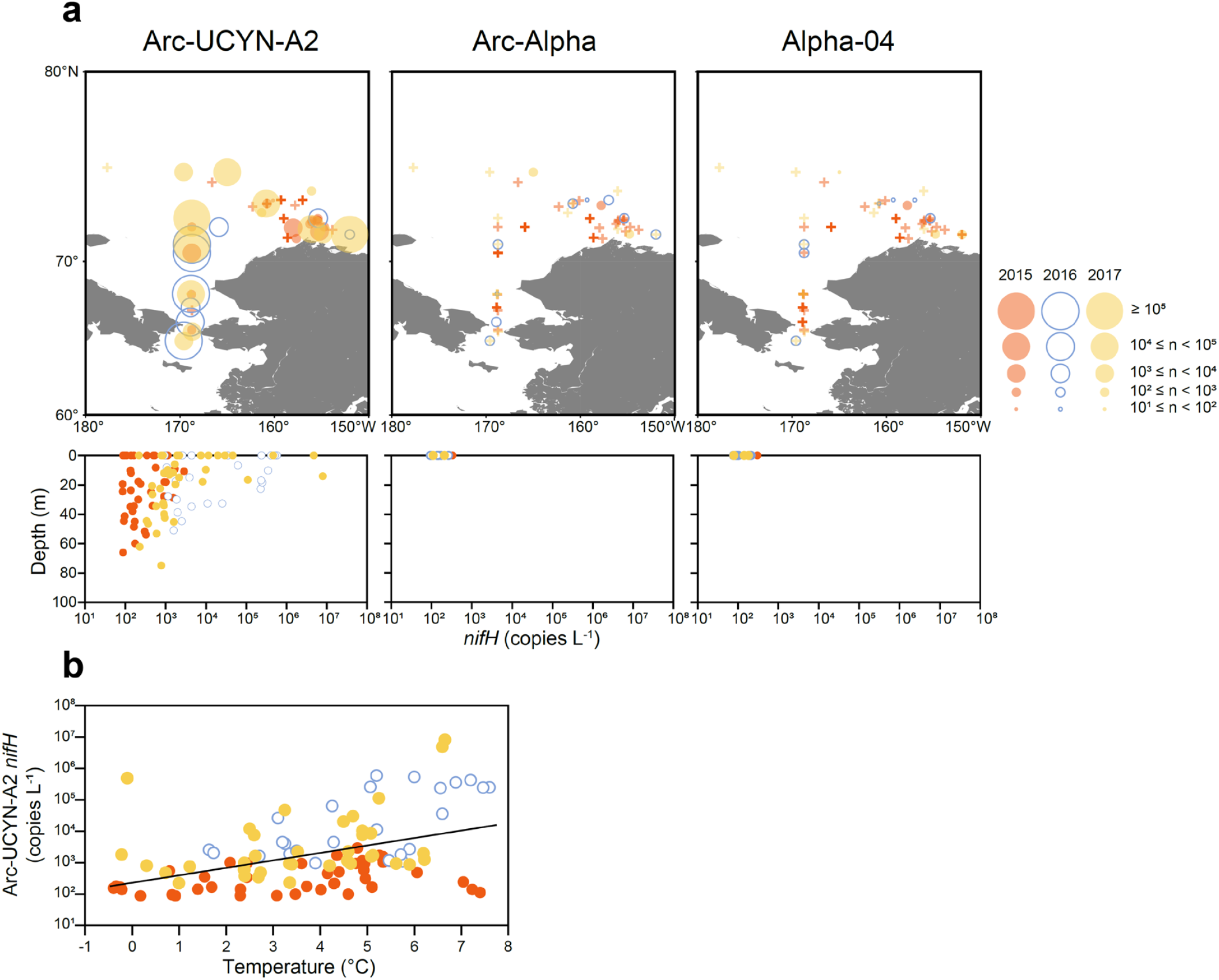
*nifH* abundance of Arctic diazotroph MAGs in the Arctic Ocean. a. Spatial and vertical distribution of *nifH* abundance. The *nifH* abundance of the spatial distribution is the maximum value at each station. b. Relationship between *nifH* abundance of Arc-UCYN-A2 and temperature. The samples were collected late summer in 2015 (orange), 2016 (blue circle), and 2017 (yellow).

## CONCLUSION

This study demonstrates the distribution and survival strategies of diazotrophs in the Arctic Ocean based on MAG information, most of which would not have been detected by a conventional PCR-based approach. Although the environments in the Arctic and tropics and subtropics are very different, we found universal diazotrophs that were distributed all over the world that have the potential for cold adaptation. In contrast, we also found Arctic-specific diazotrophs, which have specific genes that help them adapt to the Arctic environment. Considering that cyanobacterial diazotrophs occur mainly in low latitudes, the global distribution of diazotrophs follows one of three patterns: those with a low-latitude distribution, those that are Arctic specific, and those that are universally distributed.

The Arctic-occurring diazotrophs are remarkably abundant among the total metagenomic reads from the Arctic Ocean samples, indicating that they are important species in the Arctic microbial community and thus are likely to substantially contribute to biogeochemical cycles. We further showed that diazotrophs with a very small cell size (<0.2 µm) are particularly abundant in the Arctic Ocean, indicating that the standard (and even recently improved) method of using GF/F filters (pore size, 0.3–0.7 µm) ^52-54^ can markedly underestimate the rate of nitrogen fixation. The Arctic Ocean is one of the most rapidly changing oceans on earth, and nitrogen fixation in the pan-Arctic Ocean is still unknown. Our findings increase our understanding of current and possible future Arctic nitrogen fixation.

## METHODS

### Arctic Ocean metagenomes and MAG construction

The 111 metagenome data used in this study were derived from the water column of the Arctic Ocean. These samples were originally published in the Tara polar project (n = 68) ^55,56^, Polar marine reference gene catalog (n = 31) (Cao et al., 2020), and Canada Basin cruise (n = 12) ^41^ and were reanalyzed in a large-scale marine metagenome study (the OceanDNA MAG study) ^15^. The samples were collected across the entire Arctic Ocean, except for those collected during the Canada Basin cruise. Size-fractionated samples were collected during each project (0.2–3 and <0.2 µm for the Tara polar project; ≥0.2 µm for the Polar marine reference gene catalog; 0.2–3 µm for the Canada Basin cruise). The samples were collected wide depth range from 0 to 3800 m.

The OceanDNA MAG study reconstructed 52,325 qualified prokaryotic MAGs using 2,057 metagenomes derived from various marine environments ^15^. We focused on the 111 Arctic Ocean samples, which yielded 6,816 MAGs, which include 1095 species representatives. Completeness and contamination of genomes were estimated by taxon-specific sets of single-copy marker genes through the lineage-specific workflow of CheckM v1.0.13 ^57^. We explored new diazotroph genomes among the species representatives derived from the Arctic Ocean samples. The *nifH* gene was identified with a significant (<1e-05) and best hit to TIGR01287 among the TIGRFAMs HMM library ^58^ using hmmsearch (HMMER v3.3.2). The MAGs containing *nifH* were deposited in the UTokyo Repository (https://doi.org/10.15083/0002005808).

### Marine diazotroph genomes from the lower latitudes

Diazotroph genomes from low-latitude sample**s** were reconstructed from metagenomic data, which included not only existing cyanobacterial diazotrophs but also previously unknown heterotrophic bacteria ^13,14^, although genomes of *Crocosphaera subtropica* (UCYN-C) and *C. chwakensis* (formerly *Cyanothece* CCY0110) were not included. We then used these genomes as the reference genomes for the Arctic diazotroph MAGs. The genomes of *C. subtropica* and *C. chwakensis* were downloaded from the NCBI website. To perform a comparative genomics analysis of UCYN-A, we downloaded all UCYN-A genomes in the NCBI database for which the collection location could be identified.

### Taxonomic and gene annotation and metabolic and physiological potential and their clustering

All genomes used in this study were taxonomically classified using GTDB-Tk v1.7.0 (GTDB release 202) ^18^. Gene annotation was performed with hmmsearch (HMMER v3.3.2) using HMMs of Pfam ^59^, TIGRFAMs ^58^, and KOfam ^60^ databases. We further estimated the minimum generation time using Growthpred ^30^ by following the method of Royo-Llonch et al. ^29^. The lists of gene annotations for each genome were deposited in the UTokyo Repository (https://doi.org/10.15083/0002005808).

To examine the metabolic and physiological potential of each diazotroph genome, the multi-FASTA file of amino acid sequences of the genes was subjected to Genomaple™ (formerly MAPLE) ver. 2.3.2 ^61^. Genomaple™ is available through a web interface ^62^ and as a stand-alone package from Docker Hub (https://hub.docker.com/r/genomaple/genomaple). Genes were mapped to 814 functional modules defined by the KEGG ^63^, resulting in 310 pathways, 298 complexes, 167 functional sets, and 49 signatures. The module completion ratio (MCR) was calculated according to a Boolean algebra–like equation ^64^, and the Q-value was also calculated to evaluate the MCR in Genomaple™. We note that a Q-value near zero indicates a high working probability of the module ^22^. Then, we characterized the overall MCR pattern of all diazotroph genomes. The complete-linkage clustering method was used for the functional classification of diazotrophs with pairwise Euclidean distances between the overall MCR patterns for each genome using an R statistical package ver. 4.1.2 ^65^. KEGG modules with MCR values of 0% for all genomes were excluded from this analysis.

### Homology between *nifH* of Arctic diazotroph MAGs and universal *nifH* primers

The *nifH* sequence of Arctic diazotroph MAGs was tested *in silico* if it was detectable with the existing universal *nifH* primer**s** using gen_primer_match_report.py ^13^ on anvi’o ver. 7.1 ^66^.

### Phylogenetic analysis

Two types of phylogenetic tree were constructed, one for *nifH* only and one for the whole genome. The complete *nifH* sequences were aligned with MUSCLE in the MEGA11 package ^67^. Then, an *nifH* phylogenetic tree was constructed using the maximum likelihood method, and bootstrap values were determined using 100 iterations implemented in MEGA11. A whole-genome-based phylogenetic tree was constructed by PhyloPhlAn v3 ^68^, which uses ∼400 conserved marker genes. The phylogenetic tree was built with default settings and a rapid bootstrap test of 100 replicates.

### Comparative genome analysis of UCYN-A

This study used all UCYN-A genomes for which sampling locations have been clearly described (as of September 2022) (Supplementary Data 4). The method for gene annotation is detailed above. Average nucleotide identity was calculated using the ANI calculator ^69^. To visualize the UCYN-A pangenome, the anvi’o ver. 7.1 ^66^ pangenomic workflow was used. Details of the method are provided (https://merenlab.org/2016/11/08/pangenomics-v2/).

### Genome abundance of MAGs in metagenomes

We assessed the fraction of metagenomic reads recruited onto diazotroph genomes. Sequence reads of the 2,057 metagenomes used in the OceanDNA MAG study ^15^ were mapped onto 57 diazotroph genomes. Read mapping was performed with bowtie2 v2.3.5.1 ^70^ with the default setting using the quality-controlled paired-end reads of each run. If multiple sequencing runs were performed for one sample, only the run with the largest scale was used. If the sequencing run was >5 Gbps, a subset of 5 Gbps was randomly sampled. Then, the mapping results were sorted using samtools v1.9, and mapped reads with ≥95% identity, of ≥80 bp, and with ≥80% aligned fraction of the read length were extracted using msamtools bundled in MOCAT2 v2.1.3 ^57,71^. Then, the mapped reads were counted using featureCounts ^72^ bundled in Subread v2.0.0. ^73^. The genome abundance in each sample was calculated as a length-normalized count per million microbe genomes of a total community (CPMM) by the following equation,

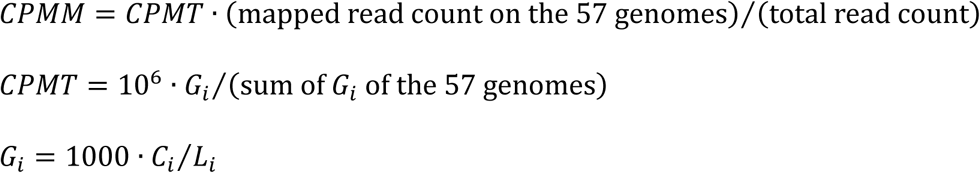

where *C*_*i*_ is a count of mapped reads on genome_*i*, and *L*_*i*_ is the length of genome_*i*. CPMT is a length-normalized count per million targeted genomes (i.e., 57 genomes). The concept of CPMT is similar to the ‘TPM’ measure, which is frequently used in transcriptome analysis ^74^.

For example, when CPMM is 1000, the genome abundance is estimated as 0.1% of the total microbe community.

### Shipboard observation and qPCR assay

We performed shipboard observations of western Arctic Ocean samples to examine the distribution of major diazotrophs in the Arctic metagenome and the environmental factors contributing to their distribution. Sampling was carried out on board the R/V Mirai MR15-03 (06 Sep to 03 Oct 2015), MR16-06 (30 Aug to 22 Sep 2016), and MR17-05C (26 Aug to 21 Sep 2017) cruises. Seawater was collected from depths corresponding to 100%, 10%, 1%, and 0.1% of surface light intensity, and from near bottom in the shelf region or from 100 m in the off-shelf region with Niskin-X bottles and a bucket. The light profiles were determined using a submersible PAR sensor just before water sampling. The depth profiles of temperature, salinity, and dissolved oxygen were measured with an SBE 911 plus CTD system (Sea-Bird Electronics). Samples for nutrient and chlorophyll *a* analyses were collected in 10-mL acrylic tubes and 290-mL dark bottles and were analyzed immediately onboard. For DNA analysis, 2 L of seawater was collected and filtered onto 0.2-◯m pore size Sterivex-GP pressure filters (Millipore). Total DNA was extracted using the ChargeSwitch Forensic DNA Purification kit (Invitrogen). Quantitative PCR (qPCR) analysis targeted the *nifH* sequences of six species (Arc-Bactero, Arc-Alpha, Arc-Gamma-01, Alpha-02, Alpha-04, and *C. chwakensis*) that were particularly highly abundant in the Arctic Ocean in the >0.2-◯m fraction and targeted *nifH* of Arc-UCYN-A2. The TaqMan probe and primer sets used here were newly designed except for Arc-UCYN-A2 (Supplementary Data 5). The qPCR analysis was conducted in triplicate using a LightCycler 480 System (Roche Applied Science, Penzberg, Germany). The r^2^ values for the standard curves ranged from 0.990 to 1.000. The efficiency of the qPCR analyses ranged from 93.8 to 100%.

## ACKNOWLEDGEMENTS

We thank all persons who contributed to the generation of the metagenomic sequence data and all persons who developed the software and databases used in this study. We also thank the captain, crew members, and participants of the R/V Mirai Arctic cruises for their cooperation at sea. We are grateful to H. Endo, Y. Nakajima, T. Masuda, and Y. Hirose for helpful discussion and support of data analyses. This study was financially supported by the Japan Society for the Promotion of Science (JSPS) KAKENHI Grant JP15H05712, JP19H04263, JP21H03583, JP22K15089, and JP22H05714, Japan Science and Technology Agency (JST) ACT-X Grant JPMJAX21BK, Arctic Challenge for Sustainability II (ArCS II) of the Ministry of Education, Culture, Sports, Science and Technology, and the FSI project “Ocean DNA: Constructing “Bio-map” of Marine Organisms using DNA Sequence Analyses” from The University of Tokyo.

