## Supplementary Information for "Distribution and survival strategies of diazotrophs in the Arctic Ocean revealed by global-scale metagenomic analysis"

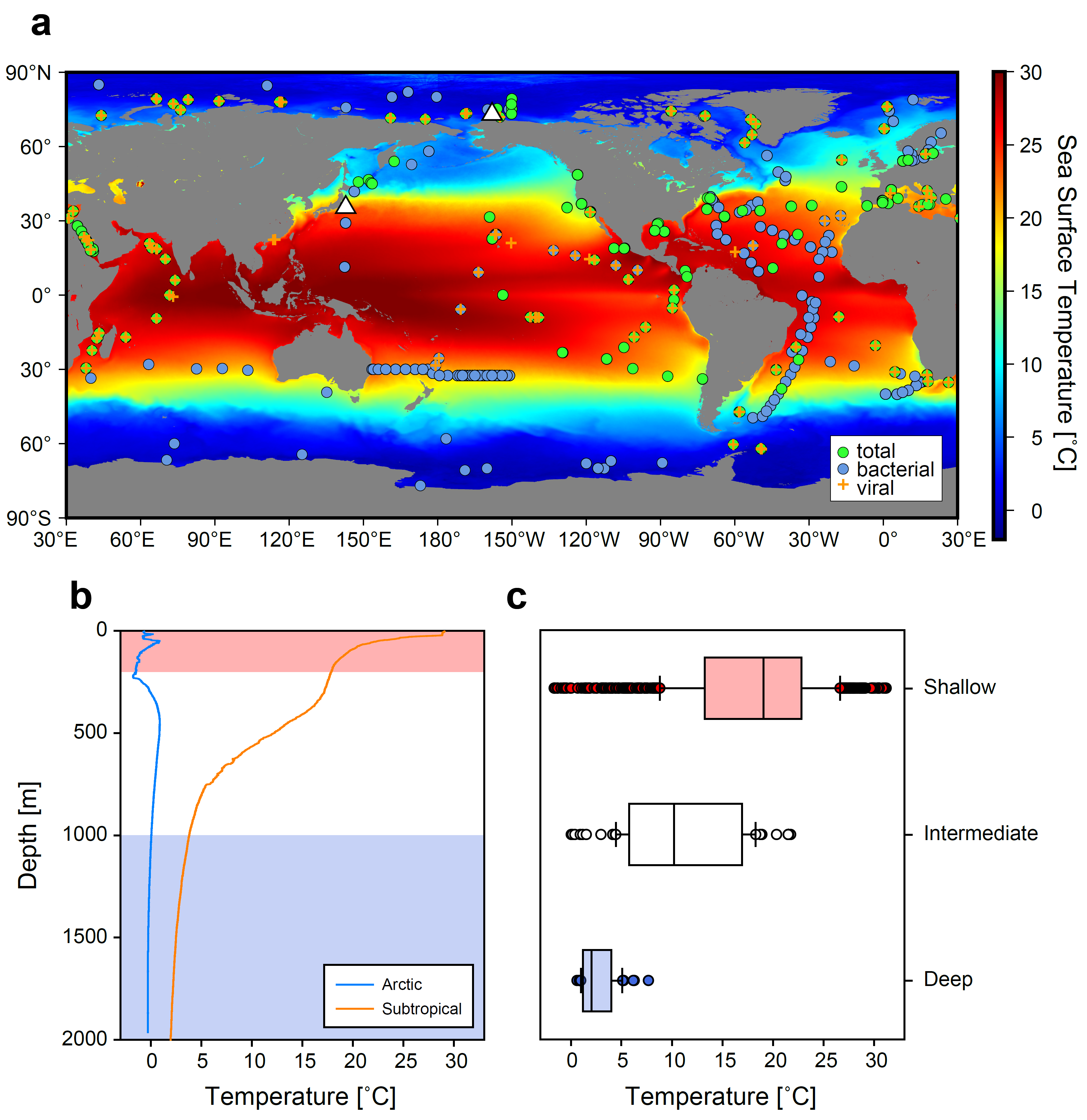

Fig. S1 Geographic distribution of the metagenomes analyzed and their water temperatures. a. Locations where samples for metagenomic analysis were collected. Green and blue circles and orange crosses indicate the locations where samples for total, bacterial, and viral fractions were collected, respectively. Background contours indicate sea surface temperature composited across the entire mission of Aqua-MODIS (https://oceancolor.gsfc.nasa.gov/). b. Temperature profile of the Arctic Ocean and subtropical ocean (white triangle in a) determined during the R/V Mirai MR20-05C cruise (https://www.jamstec.go.jp/iace/e/report/). Red, white, and blue lines indicate shallow (≤200 m), intermediate (200–1000 m), and deep (≥1000 m) layers, respectively. c. Box plot of temperature at the depths from which the metagenomic samples were collected in each layer. The line within the box indicates the median, and the upper and lower boundaries of each box indicate the 25th and 75th percentiles, respectively. The error bars indicate the 10th and 90th percentiles. Data beyond the error bars are plotted individually.

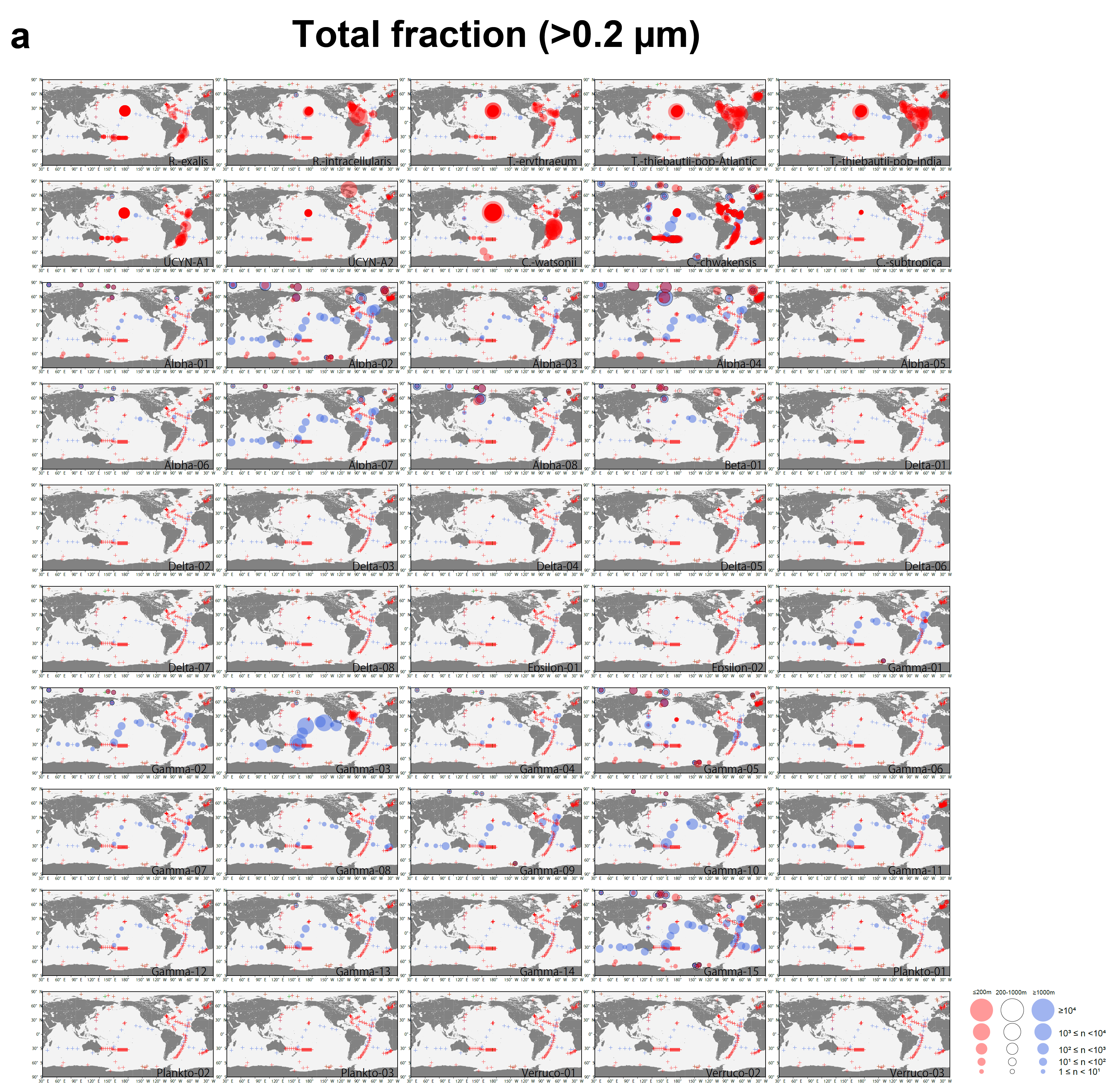

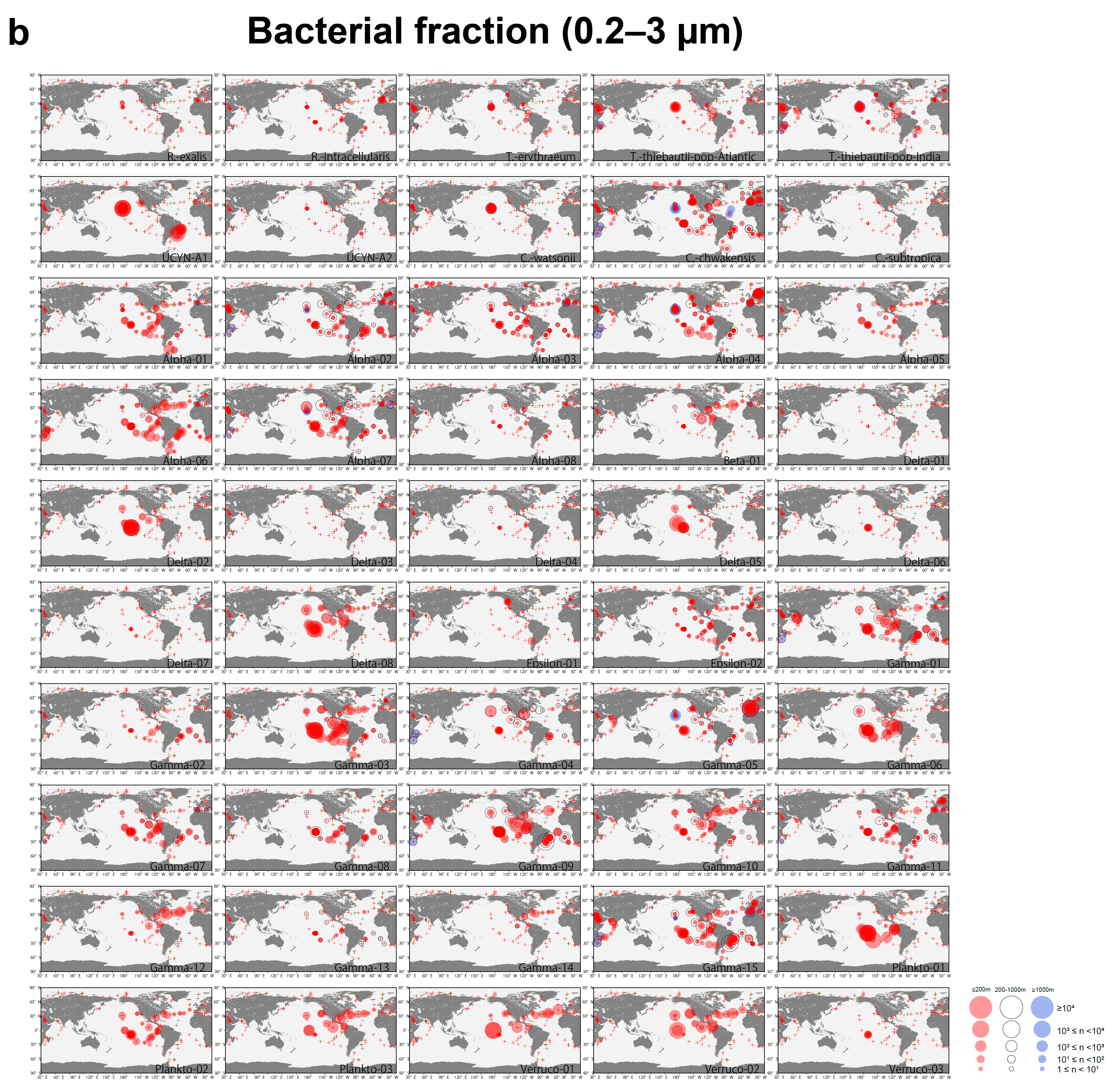

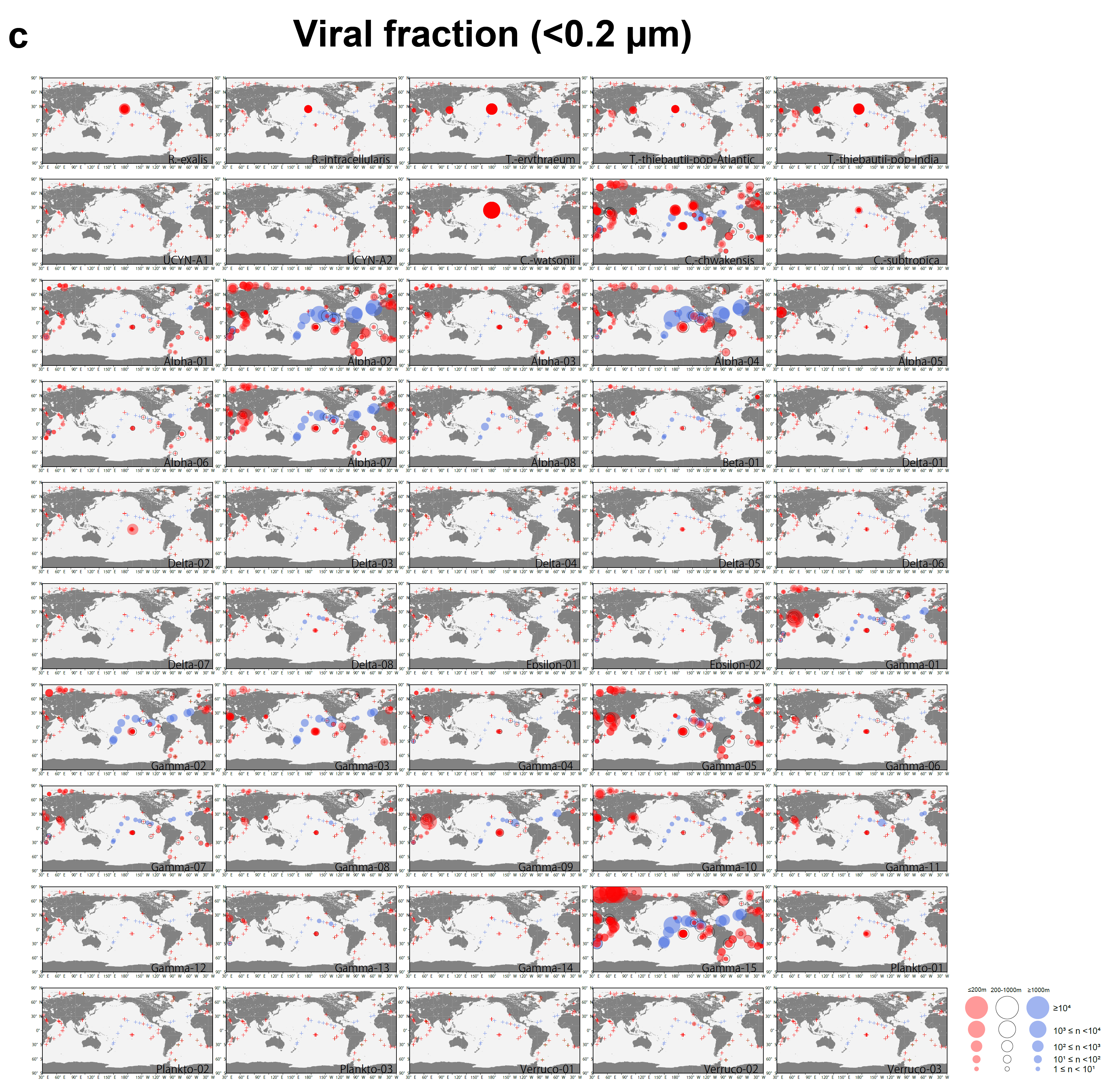

Fig. S2 Depth-resolved abundance of known marine diazotroph genomes in global ocean for each size fraction (a, total; b, bacterial; and c, viral fraction) for each layer (≤200 m, red; 200–1000 m, open circle; and ≥1000 m, blue). The area of each circle is proportional to the abundance it represents. The plus signs indicate the location with CPMM lower than 1.

Fig. S3 The pangenome of UCYN-A visualized with the anvi’o ^66^. The location from which each genome was retrieved and other details are summarized in Table S4. The dark green bar indicates genes shared by all genomes in this analysis. The ANI matrix was generated using the “anvi-compute-genome-similarity” command in anvi’o.

**Supplemental Data**

Data 1 Summary of diazotroph genomes used in this study. The table contains genome statistics, genome quality, genome-based taxonomy, minimum generation time, and number of genes that encode cold-inducible proteins and glycosyltransferases.

Data 2 The *nifH* sequence of Arctic diazotroph MAGs and its compatibility with *nifH* primers. a. The *nifH* sequence of Arctic diazotroph MAGs. b. Compatibility between *nifH* primers and *nifH* of Arctic diazotroph MAGs.

Data 3 List of the functional module completion ratio (MCR) and its Q-value as calculated by Genomaple^TM^ for each diazotroph genome.

Data 4 Summary of UCYN-A genomes used in this study. This table contains genome quality, retrieved location, reference, and *nifH* sequences.

Data 5 Primers and TaqMan probes for qPCR of *nifH*.
